# LncRNA H19 Upregulation Links Hypoplastic Left Heart Syndrome to Impaired PINK1/Parkin-Mediated Mitophagy and Ischemic Vulnerability

**DOI:** 10.64898/2025.12.16.694773

**Authors:** Xuebin Fu, Conrad L. Epting, Anshuman Sinha, Michael C. Mongé, Ming Zhao, Kristofor E. Glinton, Swetha Thirukannamangai Krishnan, My Linh Thi Nguyen, Vincent Joseph Dudley, Gregory B. Waypa, Pedro G. Lara, Tarek Kishawi, Connor W. Lantz, David Winlaw, Paul T. Schumacker, Edward B. Thorp, Zhi-Dong Ge

**Author notes:** **Address correspondence to** Dr. Zhi-Dong Ge, Cardiovascular-Thoracic Surgery & the Heart Center, Stanley Manne Children’s Research Institute, Ann & Robert H. Lurie Children’s Hospital of Chicago, Feinberg School of Medicine, Northwestern University, 303 E. Superior Stret, Chicago, IL 60611.

## Abstract

**BACKGROUND:** The myocardium in hypoplastic left heart syndrome (HLHS) exhibits immature metabolic programming, impaired mitochondrial quality control, and heightened susceptibility to ischemic and hypoxic injury during palliative surgery. The long non-coding RNA H19 suppresses translation of PTEN-induced putative kinase 1 (PINK1) mRNA and modulates mitochondrial quality control and ischemia/reperfusion injury (IRI) in adult hearts. Whether—and how—H19 regulates mitophagy and IRI in HLHS or in immature animals remains unknown.

**METHODS:** We investigated H19 regulation and its role in mitophagy and ischemia/reperfusion or hypoxia/reoxygenation injury in myocardial tissue from HLHS patients, HLHS-specific induced pluripotent stem cell–derived cardiomyocytes (HLHS-iPSC-CMs), and immature rat hearts. Mechanistic interactions among H19, PINK1/Parkin signaling, and mitophagosome formation were assessed using loss-of-function approaches.

**RESULTS:** HLHS myocardium exhibited markedly elevated H19 expression, accompanied by reduced PINK1 and Parkin protein abundance and diminished mitophagosome formation. Similar findings were observed in HLHS-iPSC-CMs exposed to hypoxia/reoxygenation and in immature rat hearts subjected to myocardial IRI. H19 knockdown in HLHS-iPSC-CMs attenuated hypoxia/reoxygenation–induced lactate dehydrogenase release and restored PINK1 and Parkin protein levels. In immature rats, myocardial H19 silencing reduced infarct size, enhanced mitochondrial PINK1 and Parkin expression, and improved post-reperfusion cardiac function for up to 28 days. Conversely, knockdown of PINK1 or Parkin reduced mitophagosome formation and exacerbated functional deterioration during IRI.

**CONCLUSIONS:** H19 upregulation impairs PINK1/Parkin-dependent mitophagy and increases susceptibility to ischemic and hypoxic injury in HLHS and the immature heart. These findings identify H19 as a key regulator of mitochondrial quality control and a potential therapeutic target for mitigating IRI in early-life cardiac disease.

Graphical Abstract

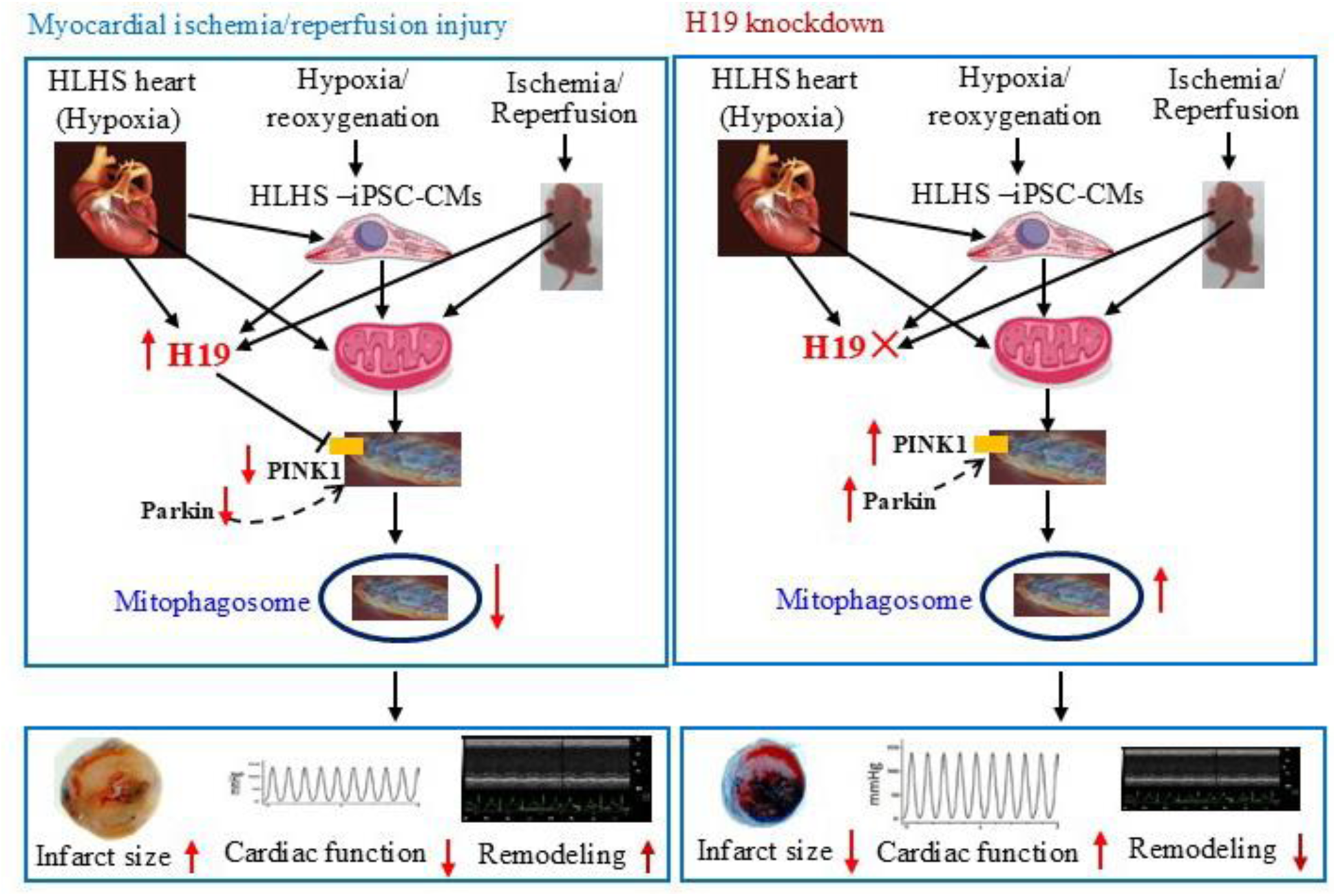

**Novelty and Significance:** *What Is Known?:*

- The myocardium in hypoplastic left heart syndrome (HLHS) exhibits immature metabolic programming, abnormal coronary perfusion, and impaired mitochondrial quality control, rendering it highly susceptible to ischemic and hypoxic injury.
- Both structural limitations and intrinsic mitochondrial dysfunction contribute to the reduced ischemic tolerance of the HLHS heart, particularly during surgical and hemodynamic stress.
- The long noncoding RNA H19 regulates mitochondrial quality control and modulates myocardial ischemia/reperfusion injury (IRI) in adult hearts.
- H19 inhibits the binding of the translation initiation factor eIF4A2 to PTEN-induced putative kinase 1 (PINK1) mRNA, thereby suppressing PINK1 protein synthesis and influencing PINK1-dependent mitophagy in adult mice.

*What New Information Does This Article Contribute?:*

- This study identifies robust upregulation of H19 in HLHS myocardium, HLHS-specific induced pluripotent stem cell–derived cardiomyocytes (HLHS-iPSC-CMs) exposed to hypoxia/reoxygenation, and in immature rats subjected to myocardial IRI.
- Elevated H19 is associated with suppressed PINK1/Parkin-dependent mitophagy, exacerbated IRI, and adverse post-injury remodeling.
- Knockdown of H19 restores mitochondrial PINK1 and Parkin protein levels, enhances mitophagy, reduces infarct size, and improves long-term recovery of cardiac function—demonstrating a previously unrecognized pathogenic role for H19 in the immature heart under stress.
- Knockdown of PINK1 or Parkin decreases mitophagosomes and exacerbates myocardial IRI in immature rats.

## INTRODUCTION

Hypoplastic left heart syndrome (HLHS) and related anomalies with a single systemic right ventricle are defined by hypoplasia of the left heart structures and ascending aorta, leading to compromised systemic cardiac output.^1^ HLHS occurs in 8–25 per 100 000 live births and accounts for up to 16% of congenital heart disease–related mortality.^2^ Current management involves a three-stage surgical palliation to restore systemic perfusion: the stage I Norwood procedure in the neonatal period, followed by the stage II Glenn and stage III Fontan procedures.^3^ During the Norwood procedure, the neonatal HLHS heart is unavoidably exposed to ischemia/reperfusion injury (IRI) resulting from aortic cross-clamping and subsequent reperfusion upon clamp release.^4^ The HLHS myocardium exhibits immature metabolic programming, abnormal coronary perfusion, and impaired mitochondrial quality control, contributing to an increased susceptibility to ischemic and hypoxic injury.^5–7^ Both structural constraints and intrinsic mitochondrial defects further reduce ischemic tolerance, particularly during surgical or hemodynamic stress,^6,8^ yet effective cardioprotective strategies for neonates remain limited.

Long non-coding RNA (lncRNA) H19 plays critical roles in cardiovascular physiology and pathology.^9–11^ It is dynamically regulated during IRI, a pathological process characterized by hypoxia-induced damage followed by oxidative stress and inflammation upon reperfusion.^12^ H19 has been implicated in regulating cell survival, apoptosis, autophagy, and inflammatory responses during IRI.^13–17^ Depending on the context and tissue type, H19 may exert either protective or detrimental effects, underscoring its complex, context-dependent role in IRI and its potential as a therapeutic target in cardiovascular disease in adults.^13,16,18,19^ However, its contribution to mitochondrial quality control and IRI in the immature heart remains unknown.

Mitochondria are central to cardiomyocyte survival, and the selective elimination of dysfunctional mitochondria through mitophagy is critical for maintaining cellular homeostasis under stress.^20,21^ The PTEN-induced putative kinase 1 (PINK1)/Parkin pathway is a well-characterized mechanism of mitophagy: PINK1 accumulates on depolarized mitochondria, recruits the E3 ligase Parkin, and promotes ubiquitination of outer mitochondrial membrane proteins to initiate clearance.^22^ In this study, we assessed the expression of cardiac H19 and PINK1/Parkin and quantified mitophagosomes (autophagosomes engulfing mitochondria) in patients with HLHS. We further examined H19 and PINK1/Parkin expression in HLHS-specific induced pluripotent stem cell-derived cardiomyocytes (HLHS-iPSC-CMs) subjected to hypoxia/reoxygenation injury (HRI). Since myocardial IRI cannot be experimentally induced in HLHS patient heart samples, we evaluated their regulation in immature rats undergoing myocardial IRI both *in vivo* and *ex vivo*. Using short hairpin RNAs (shRNA) targeting H19, PINK1, or Parkin, we investigated the effects of H19, PINK1, and Parkin knockdown (KD) on infarct size, cardiac function, mitophagosomes, and PINK1/Parkin expression in immature rats *in vivo* and ex vivo. Finally, we examined the chronic effects of H19 KD on cardiac remodeling and function following myocardial IRI. Collectively, these findings identify H19 as a critical regulator of mitophagy and myocardial IRI in HLHS and in the immature animals.

## METHODS

### Data Availability

The data that support the findings of this study are available from the corresponding author on reasonable request. A detailed description of the methods and materials can be found in the <u>Supplemental Material</u>.

### Human heart tissue collection and processing

Hear tissues were obtained from six patients with HLHS (aged 3–12 months) undergoing heart transplantation. Control tissues were collected from the donor hearts. Discarded myocardial specimens not required for clinical diagnosis were collected in collaboration with the surgical team, following protocols approved by the Institutional Review Board.

### HLHS-iPSC differentiation and transduction

HLHS-iPSC lines were purchased from Stanford University and cultured in Essential 8 medium and passaged at a 1:4 split ratio.^23^ Differentiated purified cardiomyocytes (CMs) were used in experiments.

### Vector injection in neonatal rats

Pregnant Wistar rats were purchased from Charles River Laboratory. Wistar rats aged 2–3 days were injected intracardially with 2 × 10¹² particles of lentiviral vectors carrying shRNA targeting lncRNA H19 (shRNA-H19), PINK1 (shRNA-PINK1) or Parkin (shRNA-Parkin), or corresponding scrambled controls (SCR-H19, SCR-PINK1, SCR-Parkin, respectively). Injections were performed under real-time guidance using a VisualSonics echocardiograph, as illustrated in Figure 4A.

### Myocardial IRI in immature rats

#### In vivo

Wistar rats aged 14–15 days were subjected to LAD occlusion for 60 minutes followed by reperfusion for 24 hours, as described.^24^ Control animals underwent identical surgical procedures without LAD ligation. Infarct size was assessed by coronary perfusion with 2,3,5-triphenyltetrazolium chloride (TTC) via the aortic root. The area at risk was delineated by perfusion of phthalocyanine blue dye into the aortic root following re-ligation at the LAD occlusion site.

#### Ex vivo

Rat hearts were mounted on a Langendorff apparatus for retrograde perfusion through the aorta at a constant pressure of 80 mmHg with Krebs–Henseleit buffer maintained at 37 °C, as previously described.^25^ All hearts were stabilized for 30 min, then subjected to 60 min of no-flow global ischemia followed by 120 minutes of reperfusion.

### Transthoracic Echocardiography

Non-invasive transthoracic echocardiography was performed using a VisualSonics Vevo 3100 high-resolution imaging system, as described.^26^

### Quantification of mitophagosomes in myocardium

Ultrathin sections (60–70 nm) of ventricular tissues from humans or neonatal rats were cut using diamond knives and mounted on copper mesh grids. Sections were contrasted with uranyl acetate and lead citrate and examined with a Philips CM120 transmission electron microscope operating at 60 kV, as described.^27^ Mitophagy, a critical pathway for removing damaged or dysfunctional mitochondria to maintain a healthy mitochondrial network, was assessed by identifying and quantifying double-membrane autophagosomes engulfing mitochondria (mitophagosomes).^28^

### Measurements of H19 and PINK1 and Parkin mRNA

One microgram of total RNA from each sample was reverse-transcribed into complementary DNA using the miScript Reverse Transcriptase Mix, Nucleics Mix, and HiFlex Buffer. To assess H19 expression, a 25 μL reaction mix containing cDNA (4.5 ng/well), RNase-free water, miScript SYBR Green, and primers (H19 or the housekeeping gene Rnu-6) was prepared according to the manufacturer’s instructions. The primers used were: human H19 forward 5’-TGCTGCACTTTACAACCACTG-3’, reverse 5’-ATGGTCTCTTTGATGTTGGGC-3’; rat H19 forward 5’-AAGAGCTCGGACTGGAGACT-3’, reverse 5’-GACCACACCTGTCATCCTCG-3’. Similar methods were used to determine the levels of PINK1 and Parkin mRNA.

### Western blot

Right ventricular tissues from human hearts, LV tissue from the area-at-risk of rat hearts, and right ventricular tissues from Langendorff-perfused rat hearts were homogenized in a buffer containing 3-(N-morpholino)propanesulfonic acid, EGTA, EDTA, protease inhibitor cocktail, phosphatase inhibitor cocktail, and 0.5% Nonidet™ P-40 detergent. HLHS-iPSC-CMs were lysed in ice-cold RIPA buffer. Immunoblotting was performed using standard techniques.^29^

### Measurement of lactase dehydrogenase

Lactate dehydrogenase (LDH) activity was measured using the LDH Assay Kit.

### Statistics

Comparisons between two groups were performed using Student’s *t*-test or two-way ANOVA, while one-way ANOVA was used for comparisons among multiple groups. Statistical analyses were conducted using GraphPad Prism 10.

## RESULTS

### Upregulated cardiac H19 was associated with decreased PINK1 and Parkin proteins and mitophagosomes in HLHS

We first examined the expression of cardiac H19 in patients with HLHS. H19 levels were elevated approximately fourfold compared with controls (Figure 1A). To determine whether H19 upregulation in HLHS hearts is a general consequence of heart failure, we assessed cardiac H19 expression in patients with heart failure due to aortic stenosis, hypertrophic obstructive cardiomyopathy (HOCM), and non-ischemic dilated cardiomyopathy (DCM). In contrast, H19 expression was significantly decreased in these groups compared to the control groups (Figure S1).

**Figure 1.**
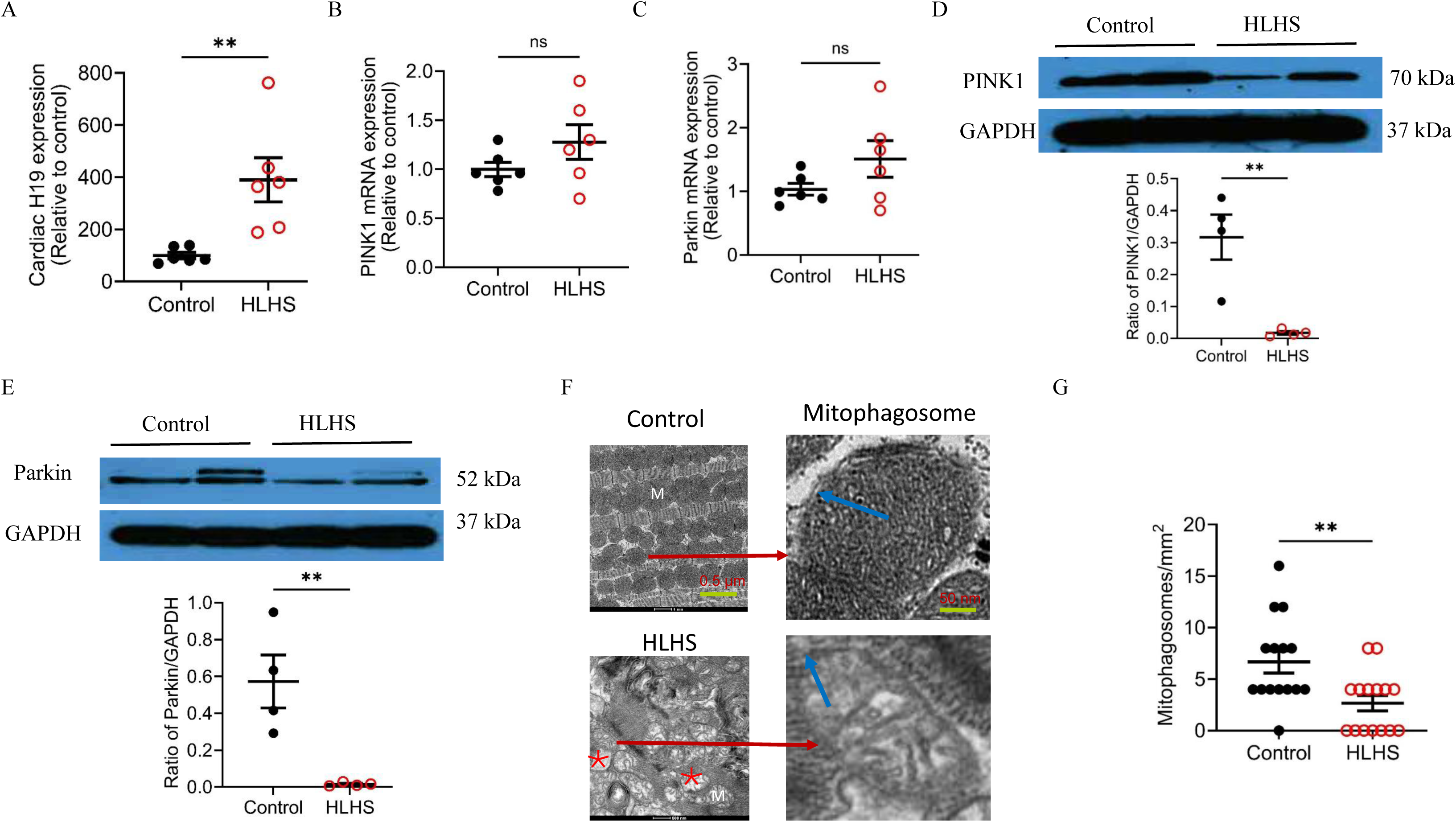
Increased myocardial H19 and reduced PINK1/Parkin protein expression and mitophagosome formation in hypoplastic left heart syndrome (HLHS). **A,** qRT-PCR analysis of cardiac H19 expression showing significant upregulation in HLHS hearts compared with controls. **B–C**, mRNA expression of PINK1 (B) and Parkin (C) showing no significant differences between HLHS and controls. **D**, PINK1 protein expression normalized to GAPDH. Top: Representative Western blots for PINK1 and GAPDH. Bottom: Quantification of PINK1/GAPDH. **E**, Parkin protein expression normalized to GAPDH. Top: Representative Western blots for Parkin and GAPDH. Bottom: Quantification of Parkin/GAPDH. **F**, Representative TEM images showing mitochondria (M) and mitophagosomes (red arrows) in control and HLHS hearts. Blue arrows indicate the double membrane structure of mitophagosomes. Mitochondrial fragmentation, disrupted cristae, and vacuolar degeneration (red asterisk) were evident in HLHS. Scale bars: left, 1 μm; right, 50 nm. **G**, Quantification of mitophagosomes per unit area demonstrating a significant reduction in HLHS hearts compared with controls. Data are presented as individual values with mean ± SEM. Student’s t-test was used for group comparisons. *P < 0.05, **P < 0.01 vs. control; ns, not significant.

A previous study has reported that H19 hinders the binding of the eukaryotic translation initiation factor eIF4A2 to PINK1 mRNA, which prevents the initiation of protein synthesis from that mRNA, thereby controlling the amount of PINK1 protein in the cell.^30^ Accordingly, we measured PINK1 and Parkin mRNA levels in HLHS hearts by the quantitative real-time reverse transcription polymerase chain reaction (qRT-PCR). Both were not significantly elevated compared with controls (P > 0.05, n = 6 hearts/group) (Figures 1B and 1C), but Western blot analysis found that the protein levels of both PINK1 and Parkin were markedly decreased in HLHS hearts (n = 4 hearts/group) (Figures 1D and 1E). PINK1 is the damage sensor, probing the integrity of the mitochondrial import pathway, and activating Parkin when import is blocked.^31^ Parkin is the effector, selectively marking damaged mitochondria with ubiquitin for mitophagy and other quality-control processes.^22^ To assess mitochondrial integrity, TEM was performed to examine myocardial ultrastructure, focusing on mitophagosomes. In controls, mitochondria and myofilaments were orderly arranged, with intact cristae and membranes. By contrast, HLHS hearts showed disorganized myofilaments and mitochondria, mitochondrial swelling, cristae dissolution, membrane rupture, vacuolization, and edema (Figure 1F). Quantification of mitophagosomes revealed a significant reduction in HLHS hearts compared with controls (Figure 1G). Together, these results indicate that cardiac H19 is upregulated while mitophagy is suppressed in HLHS.

### Upregulated H19 was associated with reduced PINK1 and Parkin proteins in HLHS-iPSC-CMs undergoing HRI

We initially attempted to induce HRI in HLHS heart tissues; however, no tissues survived after 60 minutes of hypoxia. Therefore, we performed hypoxia/reoxygenation experiments in HLHS-iPSC-CMs using 48 hours of hypoxia followed by 24 hours of reoxygenation. Compared with normoxia controls, H19 expression was significantly increased in HLHS-iPSC-CMs undergoing hypoxia/reoxygenation (HRI group) (Figure 2A). To assess the role of H19, we silenced its expression using shRNA-H19, achieving an 84% KD compared with scrambled shRNA controls. We then examined the effect of H19 KD on HRI-induced cardiac injury. As shown in Figure 2B, H19 KD did not affect LDH release under normoxia but significantly reduced LDH release in HRI-treated HLHS-iPSC-CMs. We further evaluated the effects of HRI and H19 KD on PINK1 and Parkin protein levels by Western blotting. No significant differences were observed between Control and shRNA-H19 groups under normoxia. In contrast, HRI significantly decreased the ratios of PINK1/β-actin and Parkin/β-actin, which was reversed by H19 KD (Figure 2D). These results indicate that H19 is upregulated during HRI, leading to the downregulation of PINK1 and Parkin proteins.

**Figure 2.**
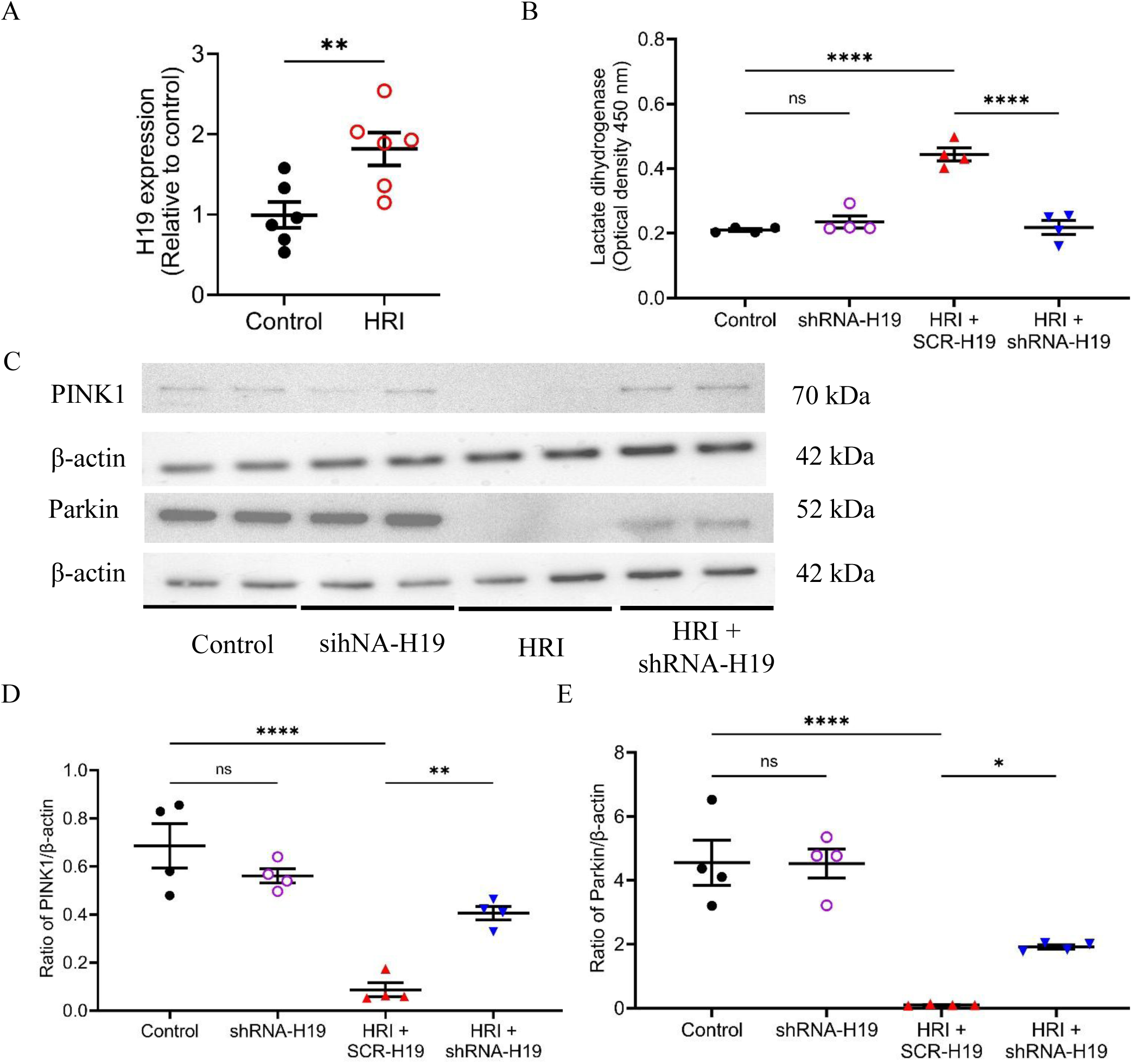
H19 upregulation during hypoxia/reoxygenation injury (HRI) suppresses PINK1/Parkin signaling in HLHS-specific iPSC-derived cardiomyocytes (HLHS-iPSC-CMs), whereas H19 knockdown alleviates cellular injury. **A**, qRT-PCR showing significant induction of H19 expression in HLHS-iPSC-CMs subjected to HRI compared with controls. **B,** H19 knockdown attenuated HRI-induced lactate dehydrogenase release in HLHS-iPSC-CMs. **C,** Representative Western blots for PINK1, Parkin, and β-actin (loading control). **D–E,** Quantification of protein expression normalized to β-actin demonstrating that HRI markedly downregulated PINK1 (D) and Parkin (E), whereas H19 knockdown partially restored their levels in HLHS-iPSC-CMs. Data are presented as individual values with mean ± SEM. One-way ANOVA was used for group comparisons. *P < 0.05, **P < 0.01, ****P < 0.001 vs. control; ns, not significant.

### Up-regulated cardiac H19 was linked with reduced PINK1, Parkin, and mitophagosomes in immature rats undergoing myocardial IRI

Previous studies have reported that H19 is regulated following IRI in adult animal hearts and other organs, but the results have been conflicting.^12,13,18,19,32–35^ We therefore investigated cardiac H19 expression and mitophagy in immature rats subjected to myocardial IRI. The myocardial ischemic time in newborns with HLHS undergoing the Norwood procedure is approximately 60 minutes.^36–37,38^ Accordingly, Wistar rats aged 14–15 days (28–32 g body weight) underwent 60 minutes of LAD coronary artery ligation, followed by 24 hours of reperfusion under anesthesia with 1.5% isoflurane (Figure 3A). Control rats underwent the same surgical procedures without LAD occlusion. Myocardial IRI induced a significant infarct size in the treated groups (n = 10 rats/group) (Figure 3B). Left ventricular ejection fraction, measured by echocardiography, was significantly reduced in the IRI group compared with sham controls (Figure 3C). Cardiac and plasma H19 levels were also significantly increased in IRI rats relative to controls (Figures 3D and S2), with reduced expression of PINK1 and Parkin proteins (Figures 3E and 3F). We then quantified mitophagosomes using TEM of left ventricular tissues. Compared with sham controls, the number of mitophagosomes was significantly decreased in the myocardium of immature rats subjected to myocardial IRI (Figure 3G). These results indicate that cardiac H19 is upregulated, while myotophagy is inhibited in immature rats subjected to HRI.

**Figure 3.**
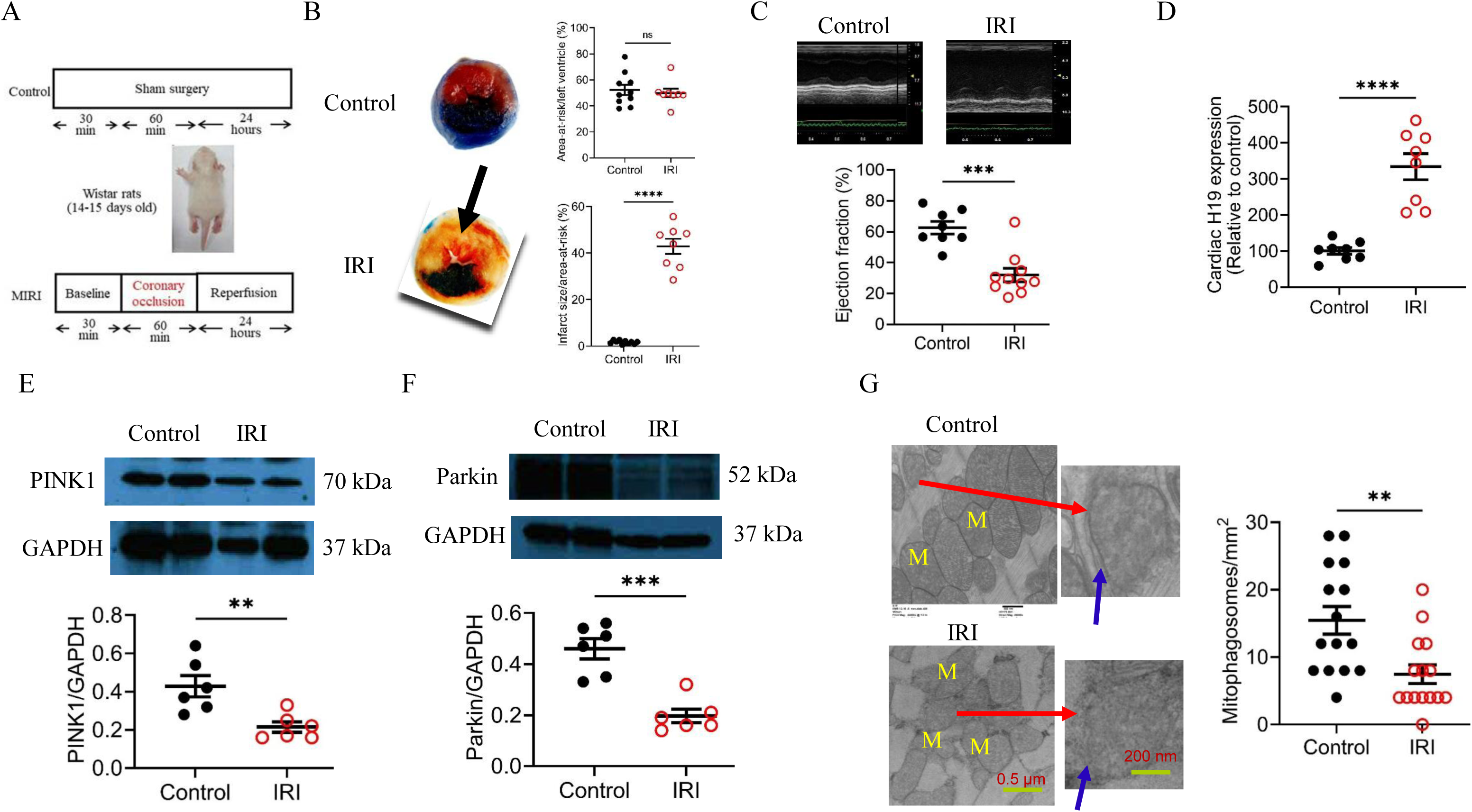
Myocardial ischemia/reperfusion injury (IRI) in immature rats induces H19 upregulation and suppresses PINK1/Parkin-mediated mitophagosome formation. **A,** Experimental protocol for sham-operated controls and IRI in 14–15-day-old Wistar rats. **B,** Area at risk (AAR) and infarct size following 60 min myocardial ischemia and 24 h reperfusion. Left: Representative TTC and phthalo blue–stained heart sections showing AAR and infarcted myocardium (arrows). Right: Quantification of AAR/LV and infarct size/AAR (n = 8–10 rats/group). **C,** Left ventricular ejection fraction determined by echocardiography (n = 8–10/group). **D,** qRT-PCR showing increased H19 expression in IRI hearts. **E,** Reduced cardiac PINK1 expression after IRI. Top: Representative Western blots for PINK1 and GAPDH. Bottom: PINK1/GAPDH ratio (n = 5 hearts/group). **F,** Reduced cardiac Parkin expression after IRI. Top: Representative Western blots for Parkin and GAPDH. Bottom: Parkin/GAPDH ratio (n = 5 hearts/group). **G,** Myocardial IRI decreased mitophagosome formation. Left: Representative TEM images showing mitochondria (M), sarcomeres, and mitophagosomes (red arrows). Blue arrows indicate the double membrane of mitophagosomes. Scale bars: left, 500 nm; right, 200 nm. Right: Quantification of mitophagosomes per unit area. Data are presented as individual values with mean ± SEM. Two-way ANOVA was used for group comparisons. **P < 0.01, ***P < 0.001, ****P < 0.0001 vs. control; ns, not significant.

### H19 KD reduced myocardial infarct size and elevated mitophagosomes in the left ventricle of immature rats

Since myocardial H19 is strongly upregulated in *in vivo* ischemic/reperfused rat hearts, we investigated whether H19 KD affects infarct size and cardiac function. Cardiac H19 was silenced in immature rats using a shRNA-H19, while a SCR-H19 served as control. Lentiviral vectors (2.0 × 10¹² genome copies) were injected into the left ventricle of 2-3-day-old Wistar rats (Figure 4A). Twelve days after injection, SCR-H19 did not alter cardiac H19 mRNA levels, whereas shRNA-H19 achieved an 83% KD compared with control (Figure 4B). Rats were then subjected to LAD occlusion for 60 minutes followed by 24 hours of reperfusion. Echocardiographic parameters prior to myocardial IRI showed no significant differences among the four groups (Table S1). The area-at-risk was comparable between shRNA-H19 and SCR-H19-treated rats; however, infarct size was significantly reduced in shRNA-H19-treated rats compared with controls and SCR-H19-treated rats (Figure 4C).

**Figure 4.**
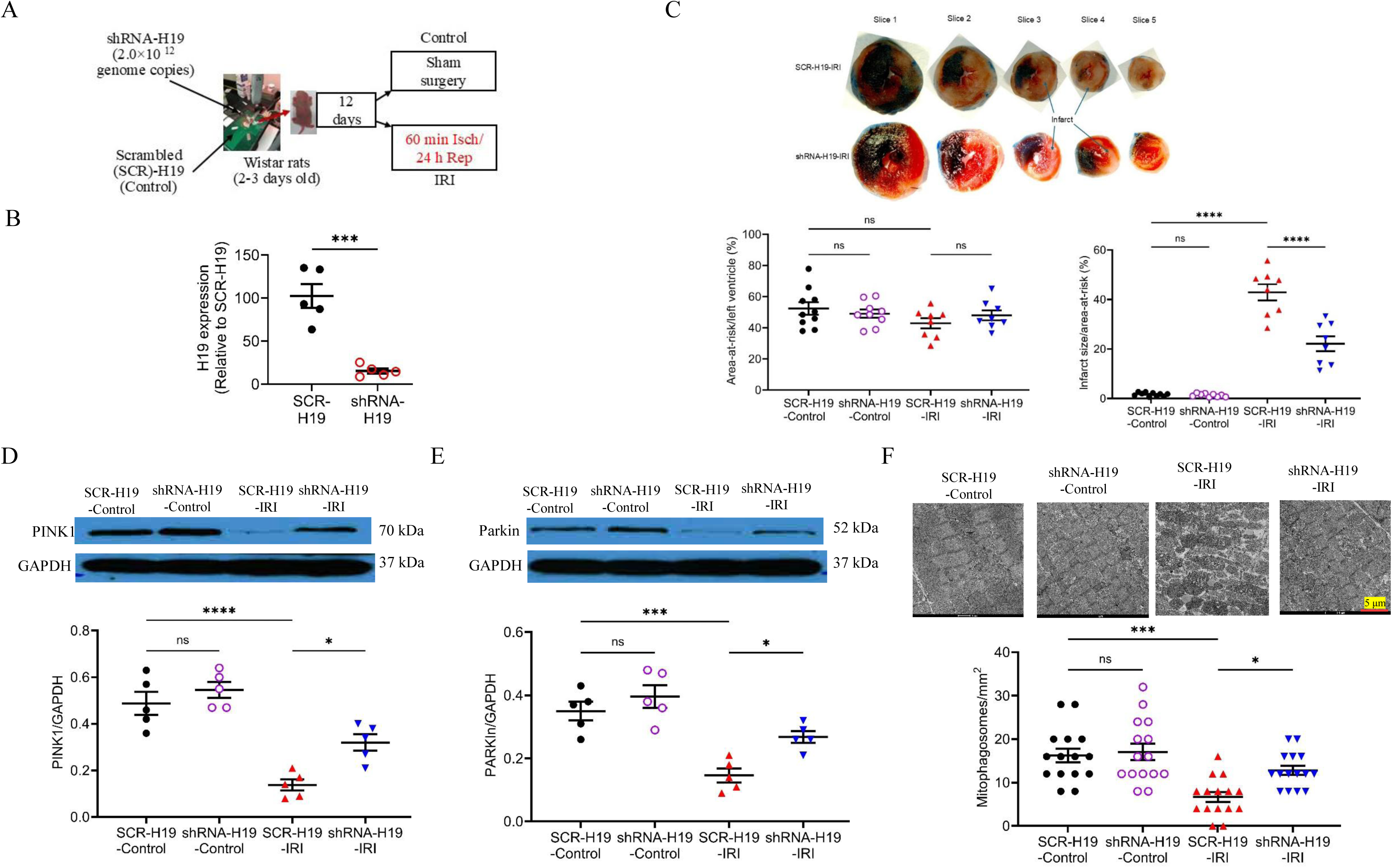
H19 knockdown attenuates myocardial infarct size and enhances PINK1/Parkin-mediated mitophagy in immature rats. **A,** Experimental protocol. Neonatal Wistar rats (2–3 days old) received intracardiac injections of shRNA-H19 or scrambled shRNA (SCR-H19). After 12 days, rats underwent sham surgery or ischemia/reperfusion injury (IRI; 60 min ischemia followed by 24 h reperfusion). **B,** Relative H19 expression in cardiac tissue assessed by qRT-PCR. ***P < 0.001 vs. SCR-H19. **C,** H19 knockdown reduced myocardial infarct size. Top: Representative TTC- and phthalocyanine blue–stained heart sections from 14–15-day-old rats subjected to IRI. Non-ischemic myocardium appears dark blue/black, viable myocardium within the area at risk (AAR) appears red, and infarcted tissue appears white. Hearts were sectioned into 5 slices from base to apex. AAR was expressed as a percentage of the LV; infarct size was expressed as a percentage of AAR. Bottom: Quantification of AAR/LV and infarct size. ****P < 0.0001 vs. SCR-H19-Control or SCR-H19-IRI. AAR/LV showed no significant differences among groups. **D,** H19 knockdown reduced cardiac PINK1 protein expression after IRI. Top: Representative Western blots for PINK1 and GAPDH. Bottom: Quantification of PINK1/GAPDH. *P < 0.05 vs. SCR-H19-IRI; ****P < 0.0001 vs. SCR-H19-Control. **E,** Representative Western blot and densitometric analysis of Parkin expression normalized to GAPDH. *P < 0.05 vs. SCR-H19-IRI; ****P < 0.0001 vs. SCR-H19-Control. **F,** H19 knockdown increased mitophagosome formation following IRI. Top: Representative TEM images of mitochondria and mitophagosomes. Scale bar: 5 μm. Bottom: Quantification of mitophagosomes. *P < 0.05 vs. SCR-H19-IRI; ***P < 0.001 vs. SCR-H19-Control; ns, not significant. Data are presented as individual values with mean ± SEM.

Mitochondrial fractions isolated from left ventricles were analyzed for the expression of PINK1 and Parkin proteins by Western blotting. The levels of PINK1 and Parkin proteins in mitochondrial fractions were not significantly altered by either myocardial IRI or H19 KD compared with sham-treated SCR-H19 rats (Figures 4D and 4E). Interestingly, shRNA-H19-treated rats subjected to myocardial IRI exhibited markedly increased mitochondrial PINK1 and Parkin proteins compared with all other groups (P < 0.05, n = 5 rats/group) (Figures 4D and 4E), indicating that H19 KD enhances mitochondrial PINK1 during myocardial IRI. Consistent with the changes in PINK1 and Parkin proteins, shRNA-H19-treated rats undergoing myocardial IRI exhibited increased mitophagosomes (Figure 4F). These results indicate that H19 regulates the expression of PINK1 and Parkin proteins and mitophagy in the left ventricle during myocardial IRI.

### H19 KD improved cardiac function and elevated mitophagosomes in the right ventricle of immature rats

To evaluate the effect of H19 KD on cardiac function, rat hearts were perfused on a Langendorff system, equilibrated for 30 minutes, subjected to 60 minutes of global ischemia, and then reperfused for 120 minutes in both the SCR-H19-IRI and shRNA-H19-IRI groups (Figure 5A). Mice in the SCR-H19-Control and shRNA-H19-Control groups underwent the same time course as the IRI groups but without global ischemia. The SCR-H19-Control and shRNA-H19-Control groups were comparable in all parameters at all time points. At baseline, during ischemia, and at 10 and 30 minutes of reperfusion, there were no significant differences in left ventricular end-diastolic pressure, developed pressure, or ±dP/dt between the SCR-H19-IRI and shRNA-H19-IRI groups (P > 0.05). However, during the 90–120-minute reperfusion period, left ventricular end-diastolic pressure was significantly lower (Figure 5B), and left ventricular developed pressure and ±dP/dt were significantly higher in the shRNA-H19-IRI group compared with the SCR-H19-IRI group (Figures 5C–5E). These results demonstrate that upregulated H19 contributes to myocardial IRI and impairs both systolic and diastolic function following reperfusion.

**Figure 5.**
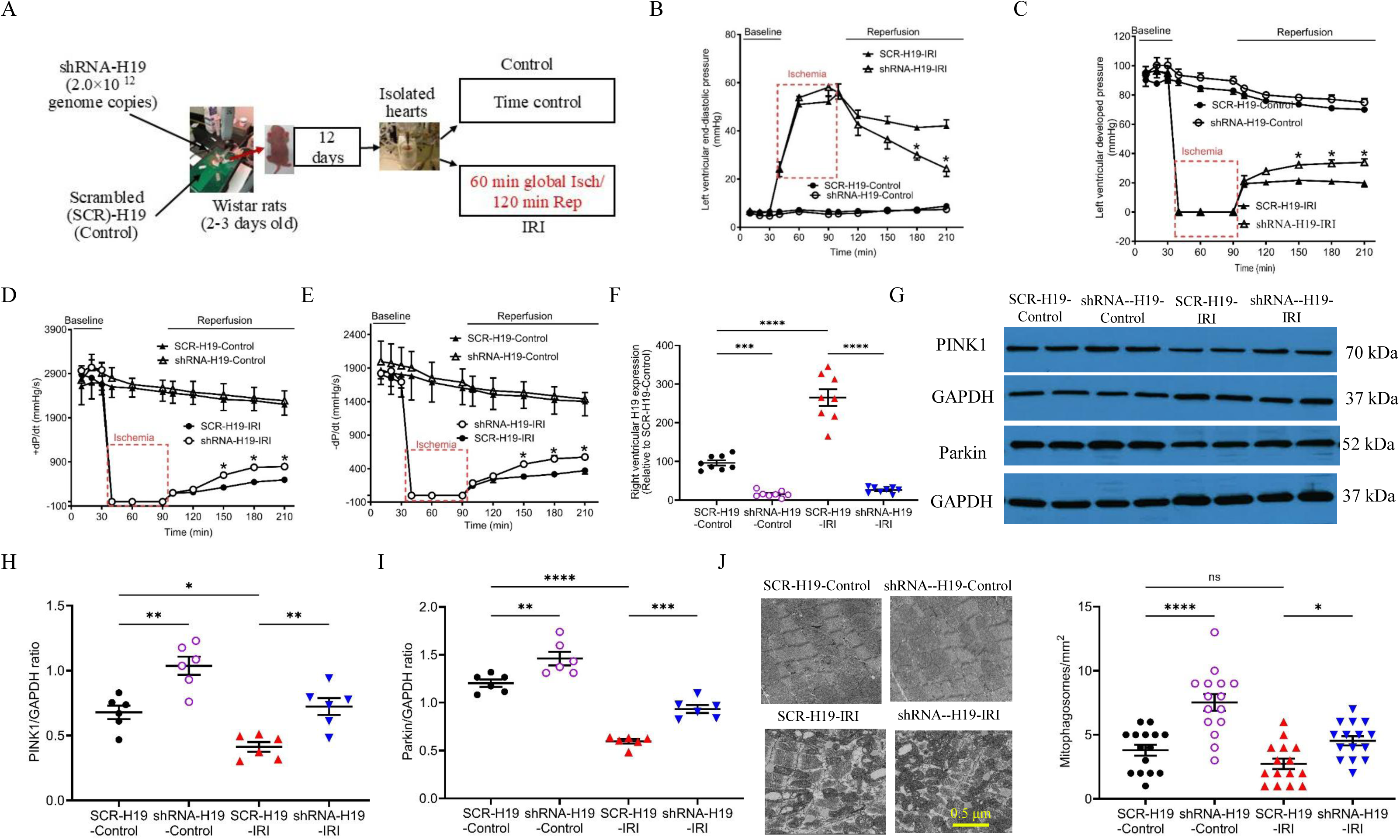
H19 knockdown improves post-ischemic cardiac function and enhances PINK1/Parkin-mediated mitophagy in Langendorff-perfused rat hearts. **A,** Experimental protocol. Neonatal Wistar rats (2–3 days old) received intracardiac injections of shRNA-H19 or scrambled shRNA (SCR-H19). After 12 days, isolated hearts were subjected to time-control perfusion or ischemia/reperfusion injury (IRI; 60 min global ischemia followed by 120 min reperfusion). **B–E,** H19 knockdown improved post-ischemic functional recovery, as indicated by reduced left ventricular end-diastolic pressure (B) and increased left ventricular developed pressure (C), +dP/dt (D), and –dP/dt (E) during reperfusion. **F,** H19 expression in right ventricular tissues of Langendorff-perfused hearts. **G–I,** Representative Western blot (G) and quantification showing that H19 knockdown increased PINK1 (H) and Parkin (I) protein levels in right ventricular tissues following IRI. GTPCH was used as a loading control. **J,** H19 knockdown increased mitophagosome formation in right ventricular tissues. Left: Representative TEM images showing mitochondria and sarcomeres. Right: Quantification of mitophagosomes. Data in **B–F** are presented as mean ± SEM and analyzed by two-way ANOVA. *P < 0.05 vs. SCR-H19-IRI. Data in **F–J** are presented as individual values with mean ± SEM and analyzed by one-way ANOVA. *P < 0.05, **P < 0.01, ***P < 0.001, ****P < 0.0001 vs. SCR-H19-Control or SCR-H19-IRI; ns, not significant.

In the human HLHS hearts, right ventricular tissues were used to assess H19, PINK1, and Parkin expression, as well as to quantify mitophagosomes. To evaluate the effect of H19 KD on PINK1 and Parkin protein expression and mitophagosome abundance in the right ventricle, we collected right ventricular tissues from Langendorff-perfused rat hearts. Compared with the SCR-H19-Control group, H19 levels were significantly downregulated in the shRNA-H19-Control group and upregulated in the SCR-H19-IRI group (Figure 5F). Notably, H19 levels were significantly lower in the shRNA-H19-IRI group than in the SCR-H19-IRI group. In contrast to the regulation of H19, the expression of PINK1 and Parkin proteins was elevated in the shRNA-H19-Control group and reduced in the SCR-H19-IRI group compared with the SCR-H19-Control group. Protein levels of PINK1 and Parkin were significantly higher in the shRNA-H19-IRI group than in the SCR-H19-IRI group (Figures 5G-5I). Consistent with these protein changes, mitophagosomes were more abundant in the shRNA-H19-Control group than the SCR-H19-Control group, and in the shRNA-H19-IRI group compared with the SCR-H19-IRI group (Figure 5J). These findings indicate that upregulated H19 suppresses mitophagy in the right ventricle of immature rats subjected to IRI.

### Either PINK1 or Parkin KD reduced mitophagosomes and exacerbated myocardial IRI

We next investigated the effects of PINK1 and Parkin KD on mitophagy and myocardial IRI. shRNA-PINK1 and shRNA-Parkin were used to knock down PINK1 and Parkin in rat hearts, respectively. KD of either PINK1 or Parkin for 12 days reduced mitophagosomes (Figures 6A and 6B), indicating that this pathway is critical for mitophagosome formation. We then examined the impact of PINK1 and Parkin KD on cardiac function after myocardial IRI. As shown in Figures 6C-6F, either PINK1 or Parkin KD impaired +dP/dt and -dP/dt at 60, 90, and 120 minutes following 60 minutes of global ischemia, suggesting that the PINK1/Parkin pathway contributes to myocardial IRI.

**Figure 6.**
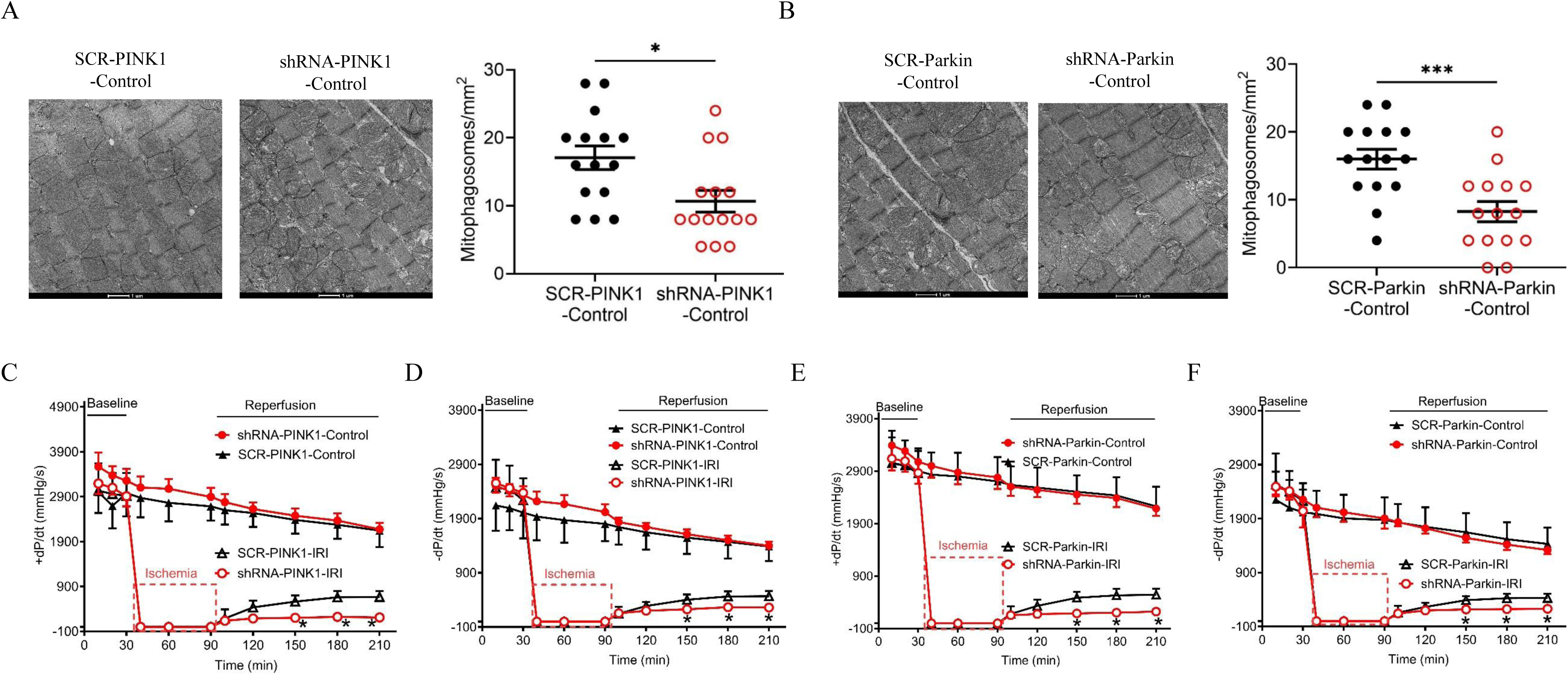
Knockdown of PINK1 or Parkin suppresses basal mitophagy and worsens post-ischemic functional recovery in isolated rat hearts. **A,** Representative TEM images (left) and quantification of mitophagosome density (right) in neonatal rat hearts transduced with scrambled control (SCR) or shRNA targeting PINK1. PINK1 knockdown significantly reduced mitophagosome number (n = 15 fields/group). Scale bar: 2 μm. *P < 0.05 vs. SCR-PINK1-Control. **B,** Representative TEM images and quantification of mitophagosome density in hearts transduced with scrambled control or shRNA targeting Parkin. Parkin knockdown markedly decreased mitophagosome number (n = 15 fields/group). Scale bar: 2 μm. ***P < 0.001 vs. SCR-Parkin-Control. **C,** Recovery of left ventricular contractility (+dP/dt) following ischemia/reperfusion injury (IRI; 60 min ischemia/120 min reperfusion) in PINK1 knockdown hearts. *P < 0.05 vs. SCR-PINK1-IRI. **D,** Recovery of left ventricular relaxation (–dP/dt) after IRI in PINK1 knockdown hearts. *P < 0.05 vs. SCR-PINK1-IRI. **E,** Recovery of +dP/dt after IRI in Parkin knockdown hearts. *P < 0.05 vs. SCR-Parkin-IRI. **F,** Recovery of –dP/dt after IRI in Parkin knockdown hearts. *P < 0.05 vs. SCR-Parkin-IRI. Data are presented as individual values with mean ± SEM.

### H19 KD improved cardiac function and prevented cardiac remodeling 28 days after IRI in rats

Following myocardial infarction, hearts undergo pathological remodeling accompanied by functional deterioration.^13,39^ We first examined the effect of H19 KD on LV systolic function over 28 days after myocardial IRI in rats. There were no significant differences in LV internal diameters at end diastole (LVIDd), end systole (LVIDs), or fractional shortening (n = 14 rats/group) between SCR-H19-Control and shRNA-H19-Control groups (Figure 7A). Compared to SCR-H19-Control rats, SCR-H19-IRI rats exhibited significantly increased LVIDd and LVIDs and decreased fractional shortening. Interestingly, shRNA-H19-IRI rats had smaller LVIDs and greater fractional shortening than SCR-H19-IRI rats, although LVIDd was not significantly different between the two IRI groups (Figure 7A). Masson’s trichrome-stained heart sections displayed significant fibrosis in the LV of both SCR-H19-IRI and siRNA-H19-IRI rats rather than in the LV of both SCR-H19-Control and siRNA-H19-Control rats. These results suggest that H19 KD improves systolic function 28 days after myocardial IRI.

**Figure 7.**
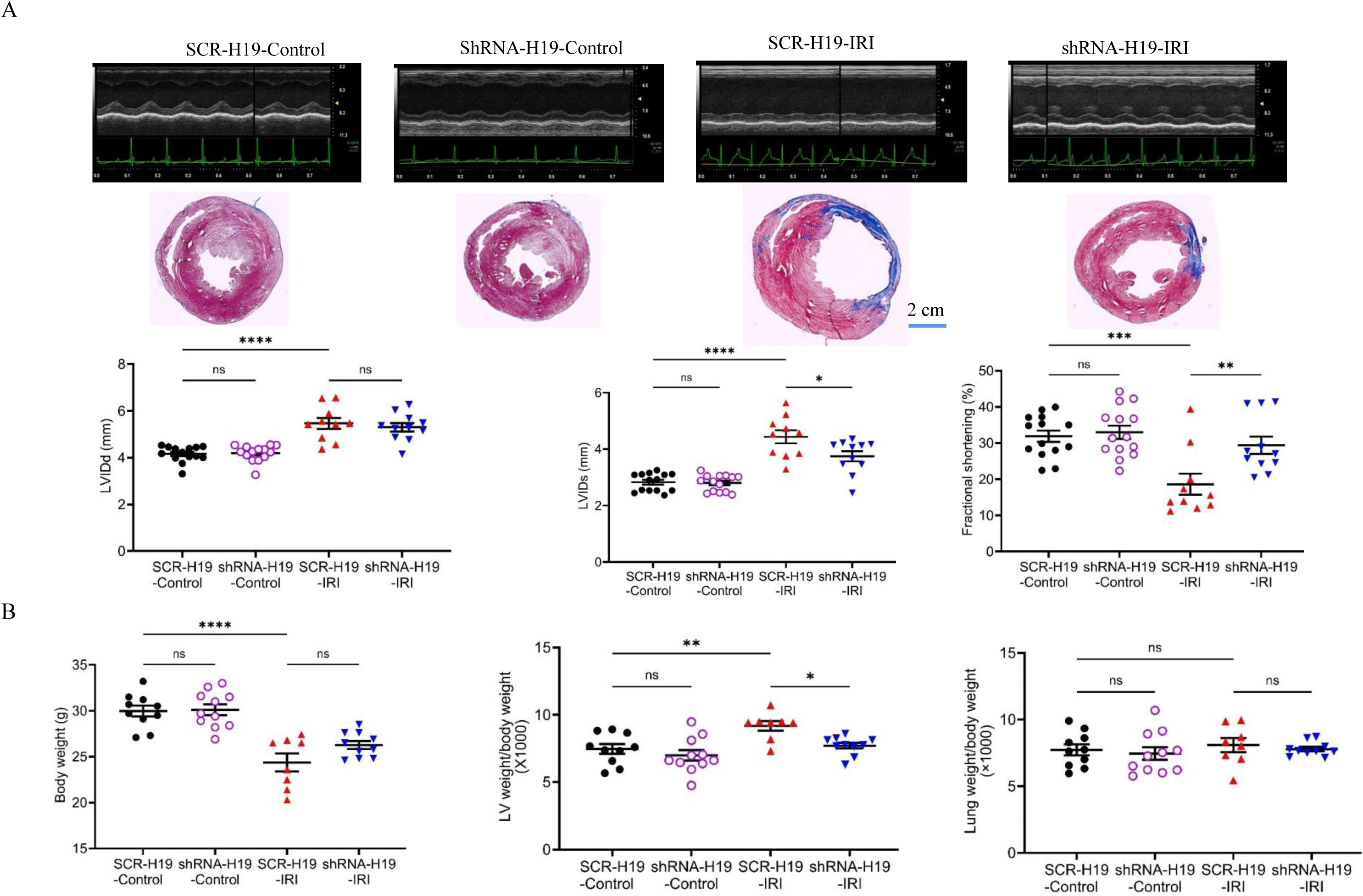
H19 knockdown improves cardiac systolic function and attenuates adverse remodeling over 28 days after myocardial ischemia/reperfusion injury (IRI) in rats. **A,** H19 knockdown improved systolic functional recovery during 28-day follow-up after IRI. Top: Representative short-axis, 2-chamber–guided M-mode echocardiographic images. Middle: Representative Masson’s trichrome–stained cross-sections at the papillary muscle level showing the left ventricle (LV) and right ventricle (RV); fibrotic tissue appears blue. Scale bar: 2 cm. Bottom: Quantification of LV internal diameter at end-diastole (LVIDd), LV internal diameter at end-systole (LVIDs), and LV fractional shortening. **B,** H19 knockdown reduced adverse remodeling after IRI. Left: Body weight. Middle: LV weight normalized to body weight. Right: Lung weight normalized to body weight. Data are presented as individual values with mean ± SEM. ns, not significant. **P < 0.01, ***P < 0.001, ****P < 0.0001 vs. SCR-H19-Control; *P < 0.05, **P < 0.01 vs. SCR-H19-IRI.

After echocardiographic assessment, rats were euthanized, and LV and lung weights were measured. Body weight and LV weight normalized to body weight were comparable between SCR-H19-Control and shRNA-H19-Control rats. Compared to SCR-H19-Control rats, SCR-H19-IRI rats had significantly lower body weight and increased LV weight normalized to body weight. Notably, LV weight normalized to body weight was lower in shRNA-H19-IRI rats than in SCR-H19-IRI rats. Lung weight normalized to body weight did not differ among the four groups (Figure 7B). These results indicate that H19 KD prevents cardiac remodeling following myocardial IRI in rats.

## DISCUSSION

The present study demonstrates that H19 is markedly upregulated in HLHS, HLHS-iPSC-CMs subjected to HRI, and immature ischemic/reperfused rat hearts, and that this elevation is associated with suppressed PINK1/Parkin-mediated mitophagy, worsened IRI, and adverse cardiac remodeling. Using patient samples, HLHS-iPSC-CMs, and immature rat models, we establish a mechanistic link between H19 dysregulation, impaired mitochondrial quality control, and cardiac dysfunction. Importantly, genetic silencing of H19 preserved mitochondrial PINK1/parkin protein levels, improved mitophagy, reduced infarct size, and enhanced cardiac functional recovery—underscoring a pathogenic role of H19 in immature hearts subjected to stress. These findings provide new mechanistic insight into the heightened vulnerability of the immature heart to ischemic stress.

Our results address an important paradox in the literature. While H19 expression is typically downregulated in adult heart failure of diverse etiologies—including aortic stenosis, hypertrophic obstructive cardiomyopathy, and non-ischemic dilated cardiomyopathy^40–42^—we show that H19 is selectively upregulated in HLHS hearts, suggesting a unique developmental or stress-response program in this congenital disease. These results are consistent with a previous study also showing down-regulated H19 in adult heart failures.^40^ One possible explanation is that immature cardiomyocytes retain a distinct metabolic and epigenetic landscape, in which H19 is activated in response to hypoxia and reperfusion stress. The divergent regulation of H19 between pediatric and adult failing hearts emphasizes the need to consider developmental stage when evaluating lncRNA-mediated mechanisms in heart disease.

Mechanistically, our data support the concept that H19 negatively regulates mitophagy through inhibition of the PINK1/Parkin pathway. Consistent with prior work linking H19 to translational control of PINK1,^30^ we observed that H19 KD restored mitochondrial PINK1 accumulation after ischemia/reperfusion, thereby facilitating mitophagosome formation and protecting against mitochondrial structural damage. Conversely, PINK1 or Parkin KD abrogated mitophagy and worsened IRI, confirming the essential role of this pathway in immature hearts. These results expand the fundings of previous studies in adults to immature hearts.^43–46^ Together, these results delineate a novel axis in which H19 acts as an upstream suppressor of PINK1/Parkin-mediated mitophagy.

Mitochondrial quality control is essential for maintaining cardiac homeostasis, particularly in the developing heart,^47^ which is characterized by a unique metabolic profile and limited adaptive capacity. Previous studies have implicated defective mitophagy in ischemic injury and heart failure.^44,45,48^ Our results extend these observations by identifying H19 as a negative regulator of PINK1/Parkin-mediated mitophagy in the immature heart. This is consistent with reports showing that lncRNAs modulate mitochondrial dynamics and cell survival under stress.^49^ By suppressing mitophagy, H19 likely promotes the accumulation of dysfunctional mitochondria, excessive oxidative stress, and cardiomyocyte death, thereby exacerbating myocardial IRI.

Functionally, H19 silencing conferred significant cardioprotection. In HLHS-iPSC-CMs, H19 KD reduced LDH release following hypoxia/reoxygenation, indicating reduced cell death. In immature rat hearts, shRNA-mediated H19 KD limited infarct size, improved post-reperfusion contractility, and enhanced both systolic and diastolic recovery. Long-term assessment further demonstrated attenuation of adverse ventricular remodeling and preservation of systolic function over 28 days. These findings extend prior reports of beneficial effects of mitophagy in adult ischemic models to immature hearts,^50,51^ highlighting that the immature heart is also highly dependent on mitophagy for stress adaptation, and that aberrant H19 expression compromises this mechanism.

Importantly, the unique pathophysiological context of HLHS magnifies the vulnerability to myocardial IRI.^52^ The right ventricle, adapted for low-pressure pulmonary circulation, must sustain systemic afterload after surgery and is therefore particularly susceptible to ischemic stress.^6^ In addition, neonatal and infant hearts have immature antioxidant defense systems and underdeveloped mitochondrial quality control, including mitophagy, further exacerbating reperfusion injury.^53^ Clinical and experimental studies suggest that inadequate protection from myocardial IRI contributes to perioperative morbidity, ventricular dysfunction, and long-term heart failure in HLHS patients.^52^ Thus, understanding the molecular basis of myocardial IRI and developing strategies to mitigate mitochondrial injury and enhance myocardial resilience are critical to improving surgical outcomes and survival in this fragile patient population.

In HLHS and immature hearts, deficiencies in mitophagy regulation likely contribute to heightened susceptibility to myocardial IRI. Neonatal cardiomyocytes exhibit underdeveloped mitochondrial dynamics and lower autophagic capacity, rendering them less efficient at clearing dysfunctional mitochondria.^54^ This impaired mitochondrial quality control may partly explain the profound ventricular dysfunction observed after palliative surgery in HLHS patients. Therapeutic strategies aimed at enhancing adaptive mitophagy—while avoiding detrimental overactivation—may therefore represent a promising approach to mitigate MIRI and improve outcomes in this vulnerable population.

In conclusion, our study identifies H19 upregulation as a key pathogenic feature of HLHS and immature ischemic hearts. By suppressing PINK1/Parkin-dependent mitophagy, H19 contributes to mitochondrial dysfunction, IRI, and maladaptive remodeling. Therapeutic targeting of H19 may thus represent a promising strategy to enhance mitochondrial quality control and improve outcomes in infants with HLHS or other pediatric cardiac conditions characterized by ischemic stress.

## LIMITATIONS OF THIS STUDY

The current results should be interpreted within the constraints of several potential limitations. First, although we demonstrate a robust effect of H19 on PINK1/Parkin-mediated mitophagy, additional downstream targets may also contribute to the observed phenotype, including interactions with other mitochondrial quality control pathways or apoptosis regulators. Second, while HLHS-iPSC-CMs provide a valuable model for HLHS, they may not fully recapitulate the complex *in vivo* environment of the developing heart. Finally, translational relevance to human patients will require validation in larger cohorts and assessment of whether H19 modulation can be safely and effectively achieved *in vivo*.

## ARTICLE INFORMATION

### Author Contributions

Z.D. Ge, E.B. Thorp, P.T. Schumacher, and D. Winlaw conceived and designed the experiments. X. Fu, C.L. Epting, A. Sinha, M.C. Monge, M, Zhao, K.E. Glinton, S.T. Krishnan, M.L.T. Nguyen, V.J. Dudley, G.B. Waypa, P. Lara, T. Kishawi, C.W. Lantz, and Z.D. Ge performed the experiments. X. Fu, C.L. Epting, A. Sinha, M. Zhao, K.E. Glinton, S.T. Krishnan, M.L.T. Nguyen, V.J. Dudley, G.B. Waypa, P. Lara, T. Kishawi, C.W. Lantz, and Z.D. Ge analyzed the data. Z.D. Ge wrote the article. Z.D. Ge, E.B. Thorp, P.T. Schumacher, and D. Winlaw proofread the article and data. All authors read and approved the article.

### Footnote

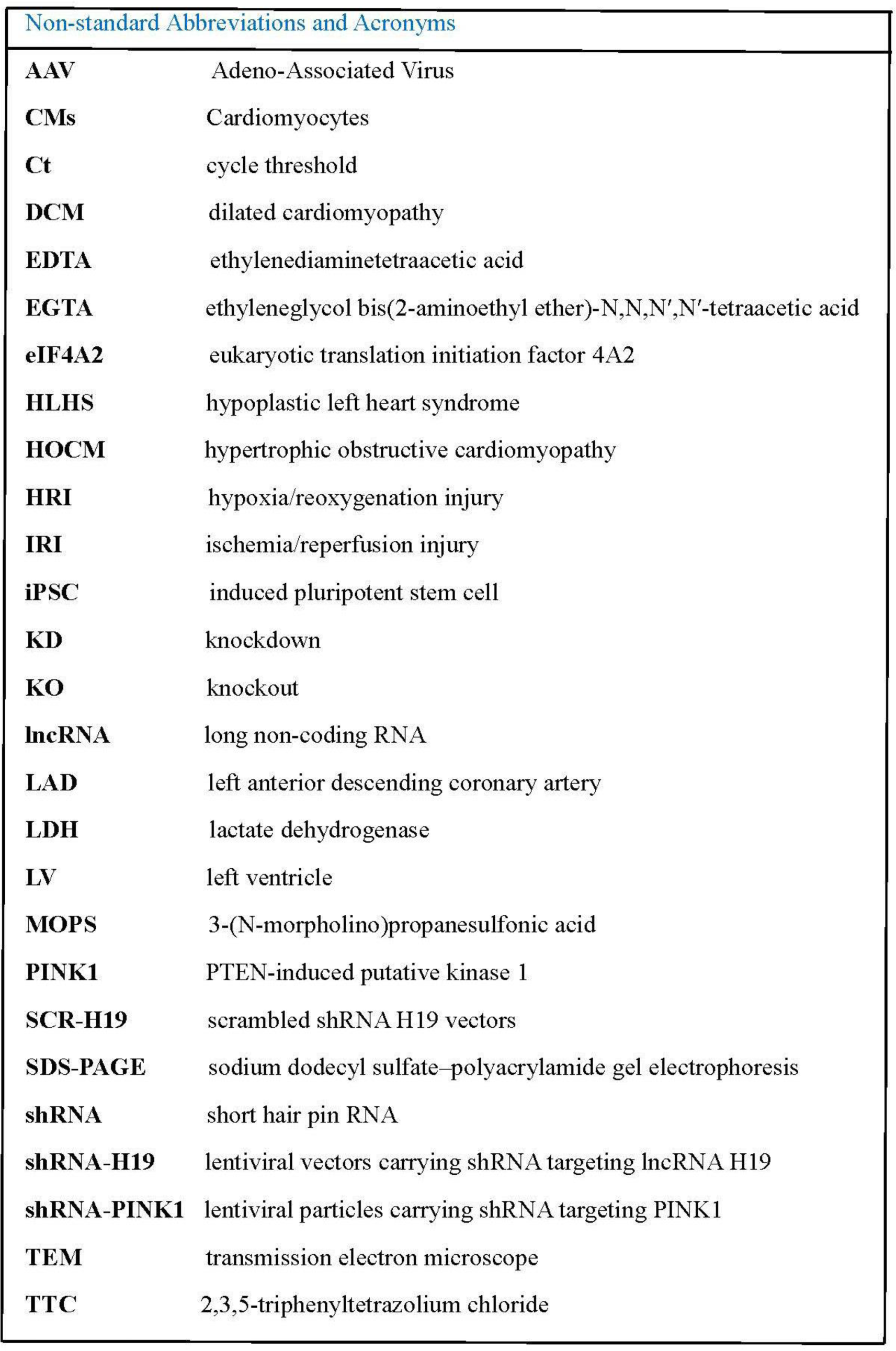

### Supplemental Material

Supplemental Materials and Methods

Tables S1–S2

Figures S1–S4

References 55–64

## FUNDING

This work was supported, in part, by two research grants 931252 and FC930151 (to Dr. Ge) from the Department of Surgery at Ann & Robert H. Lurie Children’s Hospital of Chicago, a grant from Project Bubaloo foundation (to Dr. Ge), USA, National Institutes of Health research grants R01 HL122309 (to Dr. Thorp), R35 HL177401 (to Dr. Thorp), R01 HL152712 (to Dr. Zhao) from the United States Public Health Services, Bethesda, Maryland, USA.

## Disclosures

None.

## Conflict of Interest

The authors declared no conflict of interest.

## Acknowledgements

The authors appreciate the Biorepository of Ann & Robert H. Lurie children’s Hospital of Chicago for providing human heart tissues for research.

## Notes

### Competing Interest Statement

The authors have declared no competing interest.

